# Endogenous opioid release following orgasm in man: A combined PET-fMRI study

**DOI:** 10.1101/2022.12.21.521382

**Authors:** Patrick Jern, Jinglu Chen, Jouni Tuisku, Tiina Saanijoki, Jussi Hirvonen, Lasse Lukkarinen, Sandra Manninen, Semi Helin, Vesa Putkinen, Lauri Nummenmaa

**Affiliations:** Department of Psychology, Åbo Akademi University, Turku, Finland; Turku PET Centre and Turku University Hospital, University of Turku, Turku, Finland; Department of Radiology, University of Turku, Turku, Finland; Turku Institute for Advanced Study, University of Turku, Turku, Finland; Department of Psychology, University of Turku, Turku, Finland

**Keywords:** Orgasm, Handjob, Arousal, Opioids, fMRI, Positron Emission Tomography

## Abstract

Sex is one of the most rewarding and motivating behaviours for humans. Endogenous mu-opioid receptor system (MORs) plays a key role in the mammalian reward circuit. Both human and animal experiments suggest the involvements of MORs in human sexual pleasure, yet this hypothesis currently lacks in vivo support. We used positron emission tomography (PET) with the radioligand [11C]carfentanil, which has high affinity for MORs to quantify endogenous opioid release following orgasm in man. Subjects were scanned twice: Once immediately after reaching an orgasm and once in a baseline state. Haemodynamic activity was measured with functional magnetic resonance imaging during penile stimulation from partner. The PET data revealed significant opioid release in hippocampus. Haemodynamic activity in somatosensory and motor cortices as well as hippocampus and thalamus increased during penile stimulation, and thalamic activation was linearly dependent on self-reported sexual arousal. Altogether these data show that endogenous opioidergic activation in the medial temporal lobe is centrally involved in sexual arousal.

## Introduction

The endogenous opioid system plays a key role in the mammalian reward circuit (e.g., Bozarth & Wise, 1981; DiFeliceantonio & Berridge, 2016; Nummenmaa & Tuominen, 2018), and accumulating evidence links the opioid receptor (OR) system in the regulation of sexual desire and arousal (e.g., Nummenmaa et al., 2022). Opiates are potent suppressors of sexual behaviors in both humans and animals (e.g., Pfaus & Gorzalka, 1987), and particularly μ-opioid receptor (MOR) agonists decrease human sexual desire and pleasure acutely and following chronic use (Birke et al., 2019). However, the roles of OR agonists and antagonists in exciting and suppressing sexual behaviors is complex and varies between species and conditions. In addition to their effects on sexual desire in humans (*ibid*.), opioid antagonists and agonists may also promote sexual behavior. For example, naltrexone administration both stimulated quicker-than-normal ejaculations and increased copulation rate in male rats (Rodríguez-Manzo and Fernández-Guasti, 1995). Opioid agonists may also induce copulation when injected to the medial preoptic area (Hughes et al., 1990), while striatal administration does not appear to do so, at least not consistently (Le Merrer et al., 2009).

Long-term endogenous opioidergic metabolism also contributes to individual differences in human sexual behaviour. One recent positron emission tomography (PET) study showed that MOR availability in cortical and subcortical areas, most notably in the caudate nucleus, hippocampus, and the cingulate cortices was positively correlated with sex drive in human males (Nummenmaa et al., 2022). This positive correlation is however surprising given the acute effects of exogenous opiate intake, but nevertheless corresponds well with findings on observed MOR availability in similar contexts, such as romantic and affiliative bonding in humans where higher MOR availability is consistently linked with seeking pleasure from social contacts (e.g., Turtonen et al., 2021).

In addition to opioid agonist and antagonist effects on sexual behaviour, animal studies have demonstrated endogenous opioid release following sexual activity. For example, copulation releases endogenous opioid peptides in rats, especially in the medial preoptic area of the hypothalamus (e.g., Balfour, 2004; Coolen, 2004). In humans, opioid agonists may increase pleasure, and opioid abusers have described the immediate sensations following opioid administration as euphoric and orgasmic (Chessick, 1960). Accordingly, there is evidence pointing towards the contribution of OR to human sexual drive and pleasure, but direct in vivo evidence for endogenous opioid release following sexual behaviors is lacking.

### The Current Study

Here we tested for the first time the hypothesis that sexual arousal peaking in orgasm leads to endogenous opioid release in human males. We studied healthy volunteers twice with positron emission tomography using the radioligand [11C]carfentanil which has high affinity for MORs: Once immediately after receiving tactile stimulation of the penis (i.e., a handjob) from their partner leading to orgasm and ejaculation, and once in baseline state. Acute responses to sexual arousal were quantified in a separate functional MRI study during penile stimulation by partner. We show that sexual arousal and orgasm lead to increased opioid release in the medial temporal lobe. Acute haemodynamic activity during sexual stimulation is increased in limbic regions and somatosensory cortex, while responses in thalamus reflect the moment-to-moment intensity of sexual arousal.

## Methods

### Subjects

The subjects were six heterosexual males (mean age 35.4 years, range 21.6-43.2). The exclusion criteria included a history of neurological or psychiatric disorders, alcohol and substance abuse, current use of medication affecting the central nervous system and the standard MRI exclusion criteria. All subjects gave informed, written consent and were compensated for their participation in accordance with local guidelines. The ethics board of the Hospital District of Southwest Finland had approved the protocol and the study was conducted in accordance with the Declaration of Helsinki. Their partners (all female) served as confederates for the study, providing tactile penile stimulation with their hands before one of the PET scans and during fMRI. One subject’s fMRI data had to be excluded due to technical problems.

### PET experimental protocol

Subjects were scanned with PET twice: Once in a baseline state after spending approximately 30 minutes at the PET Centre and once after receiving tactile penile stimulation from their partner (a handjob) leading to an orgasm. Partnered stimulation rather than masturbation was used to avoid perfusion confounds resulting from the physical effort involved in masturbating. Participants were offered a choice of personal lubricants for brain scans involving orgasm and were requested to abstain from sexual activity for one week prior to brain imaging. The orgasm and baseline scans were done on separate days and the order of the days was counterbalanced across participants. Before the orgasm scans the subject was cannulated and the couple was led to a private room approximately 30 minutes before the PET scan. They were informed about the timing of the PET scans and the partner was instructed that they should time the subject’s orgasm as close to the PET scan as possible, the couple was also encouraged to communicate throughout the session about the subject’s sexual arousal to time the orgasm as close to the scheduled PET scan as possible. All participants successfully achieved orgasm according to protocol.

### PET data acquisition

MOR availability was measured with radioligand [^11^C]carfentanil (Eriksson & Antoni, 2015) synthesized as described previously (Kantonen et al., 2021). Radiochemical purity of the produced [^11^C]carfentanil batches was 98.4 ± 0.3 % (mean ± SD). The injected [^11^C]carfentanil radioactivity was 258 ± 6 MBq and molar radioactivity at time of injection 370 ± 215 MBq/nmol corresponding to an injected mass of 0.46 ± 0.43 µg. Subjects were instructed to fast and abstain from smoking and drinking alcohol from 10pm the night before the day of the PET scans (i.e., subjects fasted for at least 10 hours prior to PET imaging). The subjects were also advised to abstain from physical exercise on PET scan days and the day before and abstain from sexual activities for 7 days before the scan. PET imaging was carried out with Discovery 690 PET/CT scanner (GE. Healthcare, US). The tracer was administered as a single bolus via a catheter placed in subject’s antecubital vein, and radioactivity was monitored for 51 minutes. Subject’s head was strapped to the scan table to prevent excessive head movement. T1-weighted MR scans were to correct for attenuation and for anatomical reference.

### PET image processing and data analysis

PET data were preprocessed with the Magia (Karjalainen et al., 2020) toolbox (https://github.com/tkkarjal/magia), an automated neuroimage analysis pipeline developed at the Human Emotion systems Laboratory, Turku PET Centre. Magia toolbox runs on MATLAB (The MathWorks, Inc., Natick, MA, USA), and utilizes the methods from SPM12 (www.fil.ion.ucl.ac.uk/spm/) and FreeSurfer (https://surfer.nmr.mgh.harvard.edu) as well as in-house developed tools for kinetic modeling. PET images were first motion-corrected, and co-registrated T1 MR images, after which MRI was then processed with Freesurfer for anatomical parcelation. [^11^C]carfentanil uptake was quantified by non-displaceable binding potential (*BP*ND) in 21 regions (amygdala, caudate, cerebellum, dorsal anterior cingulate cortex, hippocampus, inferior temporal cortex, insula, medulla, midbrain, middle temporal cortex, nucleus accumbens, orbitofrontal cortex, pars opercularis, posterior cingulate cortex, pons, putamen, rostral anterior cingulate cortex, superior frontal gyrus, superior temporal sulcus, temporal pole, and thalamus). *BP*ND was estimated with simplified reference tissue model in regional (Lammertsma & Hume, 1996) and voxel-level (Gunn et al., 1997) by using occipital cortex as the reference region. Due to the small sample size, we focused on region-of-interest (ROI) analyses, whereas the voxel-level *BP*ND images were used solely for illustration.

Prior to calculation of voxel-level *BP*ND images the [^11^C]carfentanil PET images were smoothed using Gaussian kernel to increase signal-to-noise ratio before model fitting (FWHM = 2 mm). *BP*ND images were spatially normalized to MNI152-space and finally smoothed using a Gaussian kernel (FWHM = 6 mm). Subsequently, regional *BP*ND across the orgasm and baseline conditions were compared using paired samples t-tests.

### fMRI experimental protocol

fMRI was conducted on a separate day. During functional imaging, the subject was covered with a blanket and their partner was sitting next to the MRI bed (see **Figure 1**). Privacy was ensured by covering the MRI control room window; security camera’s FOV only contained the subjects head at the bore of the scanner. The partner was received auditory instructions for starting and stopping to stimulate their partner in ∼10-second blocks with their hand under the blankets. Other than the timing, the handjob was unstructured, and the partner was simply instructed to stimulate the subject (i.e., their partner) as according to what they would feel pleasant. The subject was given a button box that he could use for moving a cursor on the screen to indicate his current sexual arousal on a range from 0 (not at all aroused) to 100 (orgasm). The stimulation blocks were interspersed with ∼10 second rest blocks with no stimulation (randomly chosen from durations of 10 or 12 seconds, see **Figure 2**.) Orgasm was indicated by repeated presses on the panic ball. When orgasm was reached, approximately 20 additional functional volumes were taken, and EPI scanning was terminated. Researchers did not enter the fMRI room until the participants gave a signal that it was OK to do so.

**Figure 1.**
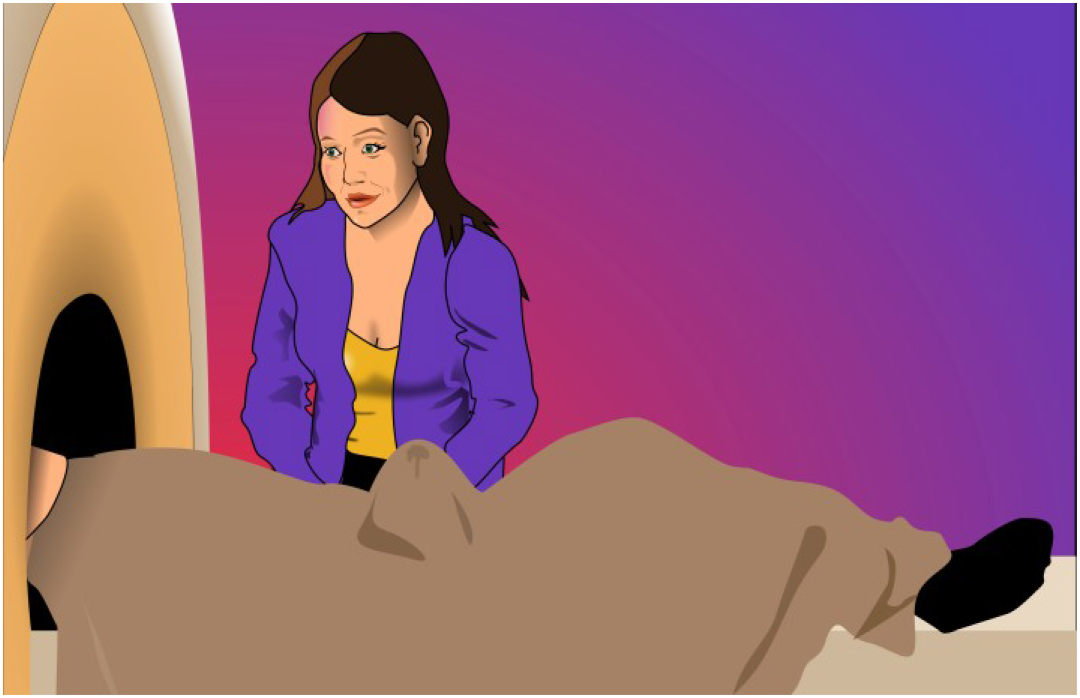
Setup for the fMRI scan and penile stimulation.

**Figure 2.**
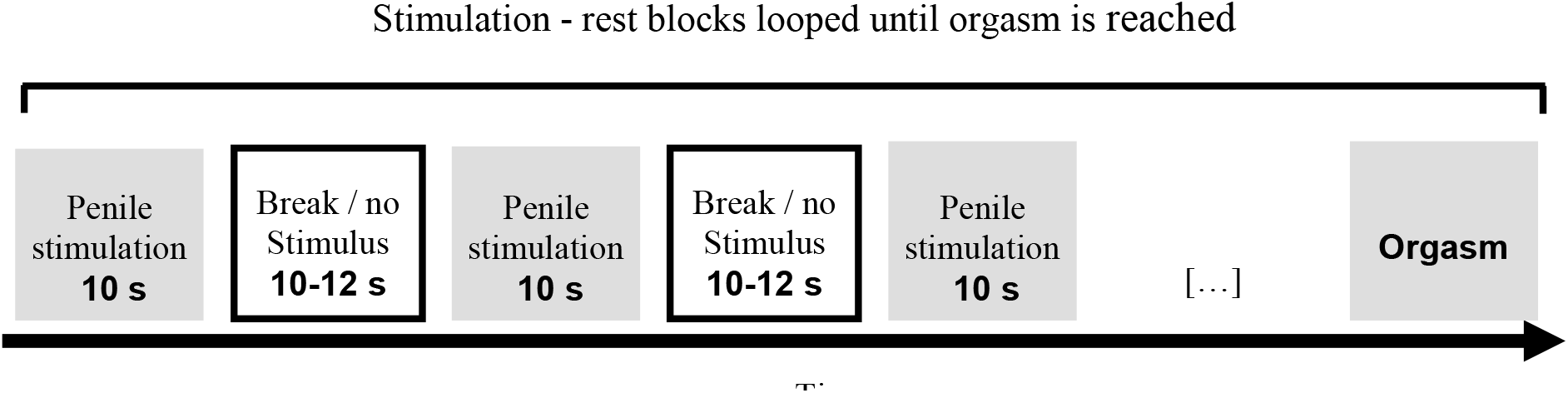
Overview of the stimulation structure in the fMRI experiment.

### MRI data acquisition and preprocessing

MR imaging was conducted at Turku PET Centre. The MRI data were acquired using a Phillips Ingenuity TF PET/MR 3-T whole-body scanner. High-resolution structural images were obtained with a T1-weighted (T1w) sequence (1 mm^3^ resolution, TR 9.8 ms, TE 4.6 ms, flip angle 7∘, 250 mm FOV, 256 × 256 reconstruction matrix). A mean of 197 functional volumes were acquired for the experiment with a T2*-weighted echo-planar imaging sequence sensitive to the blood-oxygen-level-dependent (BOLD) signal contrast (TR 2600 ms, TE 30 ms, 75∘ flip angle, 240 mm FOV, 80 × 80 reconstruction matrix, 62.5 kHz bandwidth, 3.0 mm slice thickness, 45 interleaved axial slices acquired in ascending order without gaps). A mean of 34 volumes before orgasm were discarded due to motion artefacts.

The functional imaging data were preprocessed with FMRIPREP (Esteban et al., 2019) (v1.3.0), a Nipype (Gorgolewski et al., 2011) based tool that internally uses Nilearn (Abraham et al., 2014). During the preprocessing, each T1w volume was corrected for intensity non-uniformity using N4BiasFieldCorrection (v2.1.0) (Tustison et al., 2010) and skull-stripped using antsBrainExtraction.sh (v2.1.0) using the OASIS template. Brain surfaces were reconstructed using recon-all from FreeSurfer (v6.0.1) (Dale et al., 1999), and the brain mask estimated previously was refined with a custom variation of the method to reconcile ANTs-derived and FreeSurfer-derived segmentations of the cortical grey-matter of Mindboggle (Klein et al., 2017). Spatial normalization to the ICBM 152 Nonlinear Asymmetrical template version 2009c (Fonov et al., 2009) was performed through nonlinear registration with the antsRegistration (ANTs v2.1.0) (Avants et al., 2008), using brain-extracted versions of both T1w volume and template. Brain tissue segmentation of cerebrospinal fluid, white-matter and grey-matter was performed on the brain-extracted T1w image using FAST (Zhang et al., 2001) (FSL v5.0.9).

### Regional Effects in the General Linear Model

The fMRI data analyzed in SPM12 (Wellcome Trust Center for Imaging, London, UK, (http://www.fil.ion.ucl.ac.uk/spm). To reveal regions activated by sexual stimulation, a general linear model (GLM) was fit to the data where the stimulation model (handjob vs. rest) were used as a regressor. To model the effects of sexual arousal, the moment-to-moment subject-wise sexual arousal ratings were added as parametric modulators to the model (i.e., so that the mean level of arousal per block was used as the parametric modulator). For each subject, contrast images were generated for the main effects of i) handjob and ii) sexual arousal and subjected to a second-level analysis for population level inference. Clusters surviving false discovery rate (FDR) correction (p < 0.05) are reported.

## Results

### PET

Subject-wise arousal scores for each timepoint in the orgasm and baseline scans are shown in **Figure 3**. Mean arousal rating for the orgasm was 8.1 units (scale 1-10, where 10 = extremely aroused) and the arousal following the orgasm was significantly higher than during any other point of the orgasm scan or baseline scan (*p*s < 0.05). **Figure 4** shows the mean MOR availability during the baseline and orgasm scans. The region of interest analysis (**Figure 5**) yielded statistically significant effects in hippocampus *t*(5) = 2.90, *p* = 0.03, indicating endogenous opioid release following orgasm. No other significant effects were found across the tested ROIs.

**Figure 3.**
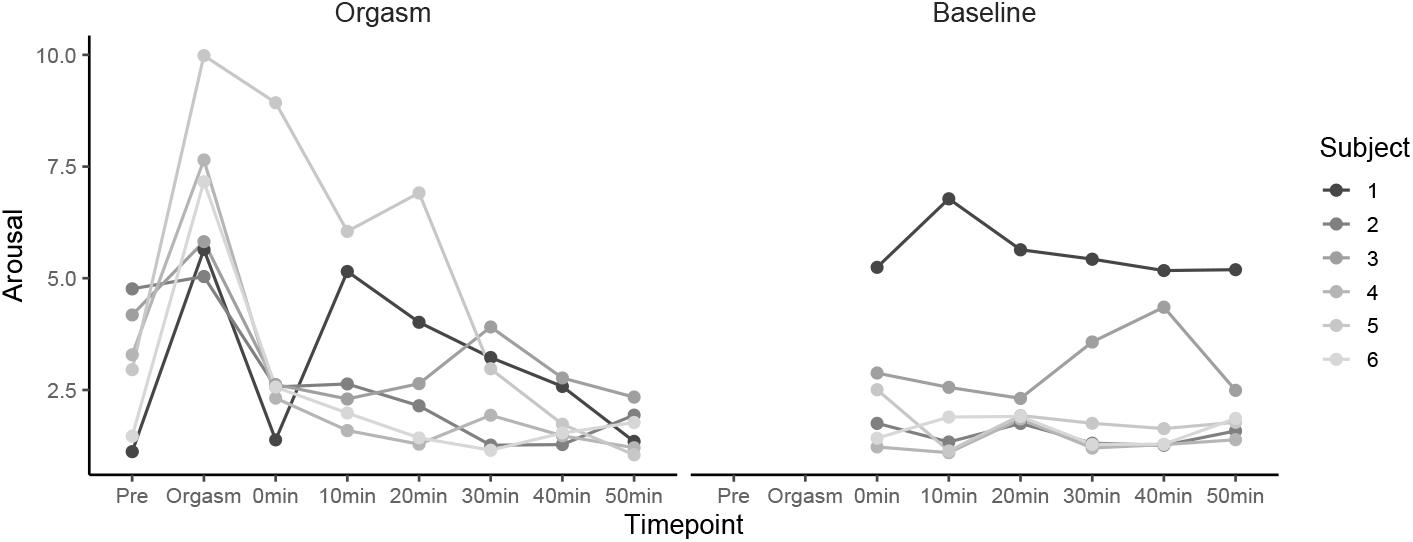
Subject-wise arousal at different timepoints during the orgasm and baseline scan. Note: The data have been jittered slightly to prevent overlap of the datapoints.

**Figure 4.**
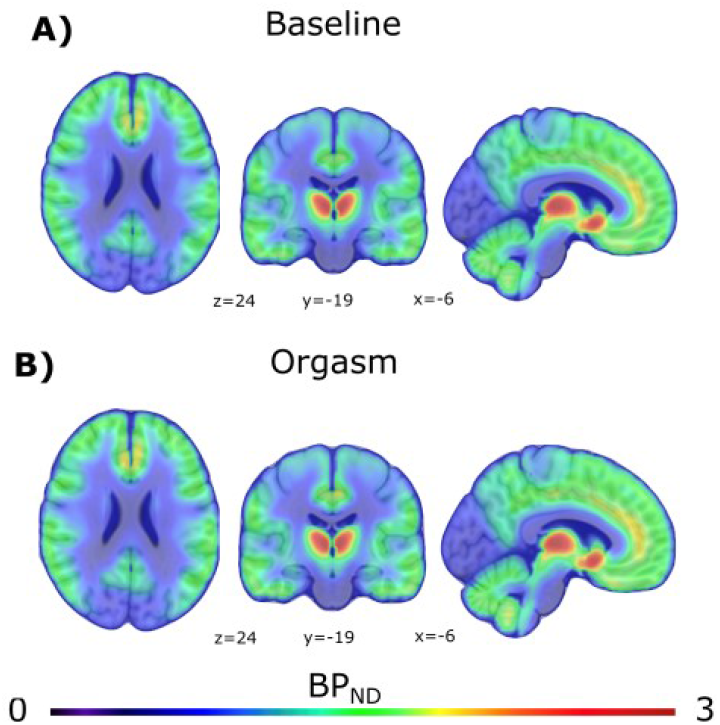
Mean MOR availability during the baseline and orgasm scans.

**Figure 5.**
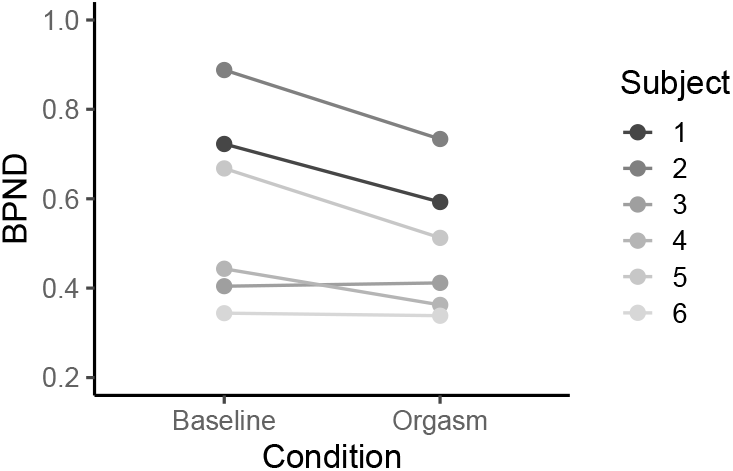
Mean subject-wise BPND in hippocampus for the baseline and orgasm scans. The difference is statistically significant at p < 0.05.

### fMRI

**Figure 6**. shows the time course of the arousal during the scans. On average it took 430 seconds for the subjects to reach orgasm and the sexual arousal increased linearly during the scans. We first assessed the haemodynamic responses to receiving a handjob minus rest. This analysis (**Figure 7A**) yielded consistent activations in the hippocampus, thalamus, anterior cingulate, posterior parietal, primary motor, and somatosensory cortices. Deactivations were observed in nucleus accumbens and insula. Finally, we modelled the parametric effects of self-reported pleasure during the scan. This analysis (**Figure 7B**) yielded significant positive effects in the thalamus and inferior parietal cortices. Negative associations were observed in insula, putamen as well as in frontal pole.

**Figure 6.**
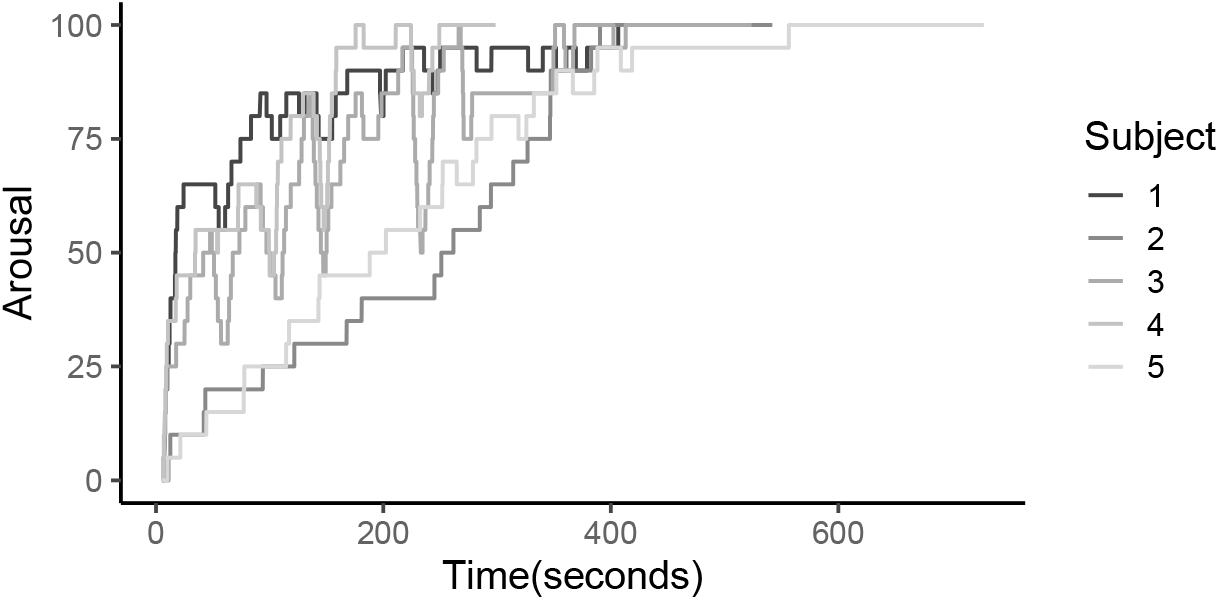
Subject-wise time courses of arousal during the scans. Lines terminate when orgasm was reached by each subject.

**Figure 7.**
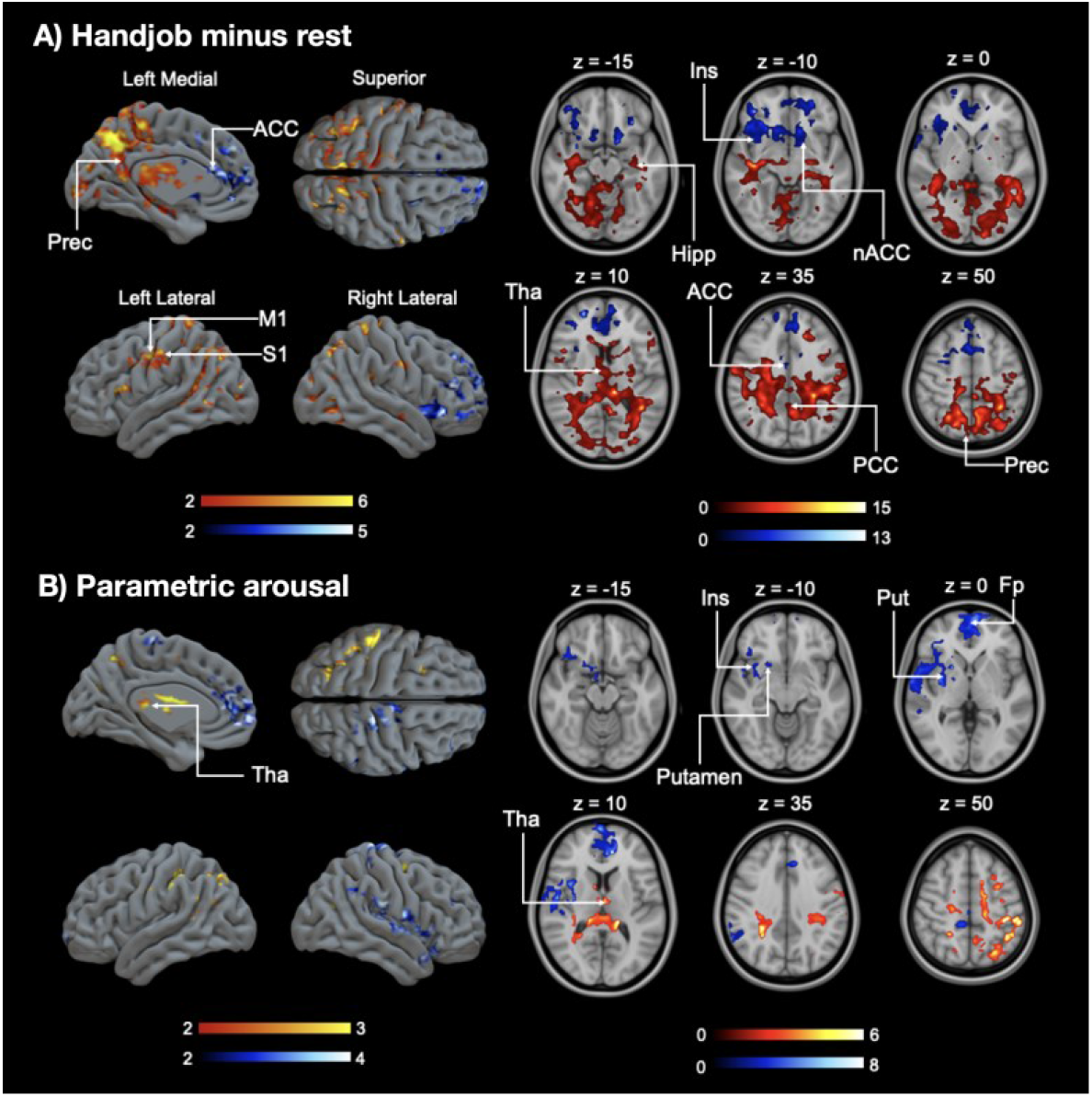
(A) Brain regions showing amplified responses during handjob minus rest. (B) Brain regions whose activity increased linearly as the function of sexual arousal during the scans. The data are thresholded at *p* < 0.05, FDR corrected at cluster level. The color bar indicates the *t* statistic range.

## Discussion

We provide, for the first time, *in vivo* evidence of endogenous opioid release following orgasm in man. PET data revealed significantly higher *BP*ND in baseline versus orgasm scan, which indicates stronger endogenous ligand binding in the orgasm scan condition. These neuromolecular changes were paralleled with robust haemodynamic responses in the hippocampus during penile stimulation. Finally, the result implied a central role of the thalamus in modulating sexual arousal. Taken together, these data yield, up to date, the most detailed picture of the functional and molecular brain basis of sexual arousal and climax in man, and support growing evidence for the general role of the endogenous opioid system in modulating the calmness—arousal axis in humans (Kantonen et al., 2020; Karjalainen et al., 2019; Nummenmaa et al., 2022).

### Opioidergic activation and sexual pleasure

The presently observed opioid release following sexual activity in human males is in general agreement with increased OR activity following sexual behaviors in other mammals (e.g., Balfour, 2004; Coolen, 2004). Furthermore, this finding accords well with human molecular imaging studies that have demonstrated endogenous opioid release following consumption of various rewards ranging from feeding to sociability (Manninen et al., 2017; Nummenmaa et al., 2018; Tuulari et al., 2017) and extends the role of the human MOR system to sexual pleasures. These data are also in line with recent PET data indicating MOR availability dependent individual differences in male sex drive, corroborating the role of ORs in modulating sexual motivation as well as sexual pleasure (Nummenmaa et al., 2022). It must nevertheless be noted due to the temporal resolution of PET we cannot conclusively state whether the presently observed opioid release reflects pleasure evoked by sexual stimulation, pleasure evoked by the orgasm or refractory activity in the post-orgasmic phase. All these options are possible given that ORs are centrally involved in hedonia, and since the predominant action of mu agonists is inhibitory in the central nervous system.

While it may seem surprising that orgasm-dependent MOR activation was only observed in hippocampus, it should be noted that i) this region is rich in MORs (Nummenmaa & Tuominen, 2018) and ii) that animal electrophysiological studies have demonstrated hippocampal activity during orgasm, evidence by high-amplitude theta activity following orgasm and ejaculation (McIntosh, Barfield, & Thomas, 1984). Furthermore, it is possible to induce theta hippocampal activity by injecting opiates into the brain stem (Leszkowicz et al., 2007). Indeed, Coria-Avila et al. (2016) suggested that the rat brain enters a “learning mode” during the post-orgasmic phase, analogous to the memory consolidation that is known to occur during sleep following learning.

It is also somewhat surprising that we did not observe significant MOR activity in the hypothalamus, given its well-described role in sexual functioning as well as the evidence from rodent studies (e.g., Hughes et al., 1990) showing copulation-inducing effects of opioidergic stimulation of the medial preoptic area. Due to the limited sample size, this may simply reflect insufficient statistical power, yet we did not observe any hypothalamic effects even at lower statistical thresholds. It must also be noted that the hypothalamus is a small structure, whose accurate quantification using PET with limited spatial resolution is complicated.

### Haemodynamic responses to sexual pleasure

Brain activity increases generally during sexual reward in both humans and animals (e.g., Yang et al., 2007), and in the present study, we observed specific activation in several brain areas during penile stimulation and orgasm. In the fMRI data analyses, we observed increased activations in the hippocampus and thalamus during penile stimulation versus rest, the former according with increased MOR activation observed in the PET study. Thalamus has general role in modulating arousal and awareness (Schiff, 2008) and the male thalamus becomes activated during penile erection acting as a “relay station” transmitting peripheral sexual sensations to the brain (see Temel, Visser-Vandewalle, Ackermans, & Beuls, 2004, for review). We also observed activation in the anterior cingulate, posterior parietal, primary motor, and somatosensory cortices, all areas of the brain that have previously been shown to become activated during sexual arousal and orgasm in heterosexual men (see Stoléru et al., 2012, for review). Surprisingly, however, we observed deactivations of the nucleus accumbens and insula in the present study. While previous studies have shown that both areas tend to become activated rather than deactivated during penile stimulation, and while previous results regarding activation of the nucleus accumbens have been somewhat ambiguous (*ibid*.), there is robust evidence for insular activation as a result of penile stimulation in humans. For example, Georgiadis and Holstege (2005) observed strong activation of the right posterior insula during penile stimulation, and similar activation patterns were observed in the majority of studies included in the review conducted by Stoléru et al. (2012). We do not have a clear-cut explanation for the presently observed deactivations, but we speculate that they may pertain to refractory responses resulting from the onset-offset blocking of the tactile stimulus.

### Limitations

We scanned only men, so the results do not necessarily translate to women. The sample size was limited due to the complex multimodal imaging setup, however significant effects were nonetheless observed in both PET and fMRI. We initially aimed at quantifying brain responses during the orgasm phase in fMRI, but ultimately motion artefacts rendered this analysis impossible despite careful fixation of subjects’ heads with pads. Future studies aiming at measuring brain activity during climax should implement even stricter prospective motion prevention such as bite bars or custom-molded face masks.

## Conclusions

In the present study, we observed endogenous opioid release in the male brain following orgasm in six healthy volunteers. The results, obtained with PET brain imaging using the μ-opioid receptor-specific ligand [^11^C]-carfentanil, were consistent with previous research on both humans and animals, however, *in vivo* evidence for endogenous post-orgasmic opioid release had not – to our knowledge – been published before. In a parallel fMRI experiment, we observed activation of above all the hippocampus. Altogether these data show that endogenous opioidergic activation in the medial temporal lobe is centrally involved in sexual arousal, while modulation of sexual arousal in thalamus and striatum may be supported by other neuromodulators.

## Acknowledgements

The study was supported by the Sigrid Juselius Stiftelse and Academy of Finland (294897, 332225). We thank Henry Karlsson for his help with generating the audio cue stimuli for the experiment, Juha Lahnakoski for sketching Figure 1, and Profs. Juha Rinne and Matti Laine for discussions on earlier versions of the research protocol.

